# A network model for patient-derived drug response in breast cancer integrating multi-omics datasets

**DOI:** 10.1101/2025.06.09.658757

**Authors:** Banabithi Bose, Barbara Stranger, Serdar Bozdag

## Abstract

The widespread availability of multi-omics tumor profiling has enabled detailed molecular characterization of individual tumors, paving the way for more effective, less toxic, and patient-specific therapies. However, widespread compound screens in human patients are constrained by ethical and logistical challenges, underscoring the need for computational models capable of predicting *in vivo* drug response. Here, we introduce *PDDRNet-MH*, a multiplex heterogeneous network-based framework that integrates genomic, transcriptomic, and epigenomic tumor profiles with drug chemical structures, pharmacological activity, and side-effect data to infer personalized drug responses. *PDDRNet-MH* constructs an integrated network connecting patients, cell lines, drugs, and genes, with each component represented by four biologically and pharmacologically informed similarity layers. This design enables the systematic propagation of drug-biomarker associations across modalities. We applied *PDDRNet-MH* to breast cancer patient data from The Cancer Genome Atlas and benchmarked its predictive performance against state-of-the-art methods on eleven FDA-approved breast cancer drugs. *PDDRNet-MH* achieved consistently high accuracy, with perfect prediction scores for gemcitabine and vinorelbine (Area Under the Receiver Operating Characteristic Curve [AUC-ROC] = 1.00; Area Under the Precision-Recall Curve [AUC-PR] = 1.00) and near-perfect scores for methotrexate and zoledronate (AUC-ROC = 0.95; AUC-PR = 0.99), demonstrating its ability to robustly distinguish sensitive from resistant patients. Biologically, *PDDRNet-MH* accurately prioritized established clinical biomarkers, including HER2 (ERBB2) for lapatinib and BRCA1/2 for doxorubicin and cyclophosphamide. Beyond known associations, the model identified additional genes within the HER2 amplicon on chromosome 17q12, including STARD3, MIEN1, and PPP1R1B, whose amplification was significantly associated with elevated drug response scores, suggesting potential roles in HER2-targeted therapy. These findings highlight the ability of *PDDRNet-MH* to recover and extend clinically relevant drug-biomarker associations, supporting its utility in guiding precision oncology.

## Introduction

Cancer remains one of the leading causes of death in the United States, with over two million new cases and more than 600,000 deaths projected annually^1^. Despite advances in detection and treatment, traditional approaches that rely on cancer type to guide therapy fail to account for the profound molecular and genetic heterogeneity of cancer. A one-size-fits-all treatment model overlooks critical differences between tumors, even within the same tissue type, leading to suboptimal outcomes and exposing many patients to severe, often avoidable, toxicities^2–5^. Mounting evidence shows that matching targeted therapies to specific molecular alterations improves outcomes, reduces adverse events, and can overcome mechanisms of drug resistance that emerge during treatment^6,7^. Therapies based on tumor molecular features, rather than tissue of origin, often better predict therapeutic response^8,9^. In light of this, precision oncology, anchored in the molecular profiling of each patient’s tumor, is not only scientifically justified but ethically imperative to improve survival, minimize harm, and deliver more effective, individualized cancer care.

A major barrier to implementing precision oncology is the challenge of characterizing drug response. While randomized clinical trials remain the gold standard for evaluating drug efficacy and toxicity, testing multiple chemotherapeutic agents directly in patients is ethically and logistically infeasible due to concerns over safety, toxicity, and trial complexity^10^. As an alternative, cancer cell lines are frequently used as proxies for patient tumors, as exemplified by large-scale drug screening efforts like the Genomics of Drug Sensitivity in Cancer (GDSC)^11^ and the Cancer Cell Line Encyclopedia (CCLE)^12^. However, prolonged culturing and genetic drift *in vitro* cause these cell lines to diverge from primary tumors, limiting their translational relevance^13,14^.

To address the limitations of current drug response approaches, computational approaches have emerged as a promising strategy for predicting therapeutic efficacy *in silico* using molecular data. Early efforts leveraged gene expression data and machine learning to infer drug sensitivities in cancer cell lines and extrapolate findings to patient samples. For example, network-based models have integrated gene expression, drug chemical structures, and protein-protein interactions to predict drug response in cell lines^15^. However, drug responses in cell lines often fail to accurately reflect those in patients^13,14,16,17^. To improve clinical relevance, novel methods such as the imputed drug-wide association study (IDWAS) framework used ridge regression to estimate drug sensitivity in cancer patients^18^, while support vector machines with recursive feature elimination (SVM-RFE) classified cancer patients as drug-sensitive or resistant based on gene expression features^19^. Despite these advances, most models overlook key genomic features, such as copy number alterations, somatic mutations, and epigenetic modifications, that are known to influence drug response^20–23^. Moreover, few existing models incorporate chemical or toxicological properties of drugs, including adverse effects, which can substantially influence outcomes^3,24^. To overcome these limitations, we developed *PDDRNet-MH*, a patient-centric drug response prediction model that integrates multi-omics data from cancer patients with drug chemical properties and side-effect profiles.

Graph-based algorithms have shown strong performance in modeling biological systems due to their ability to represent complex relationships among functionally related entities. One widely used approach is based on the principle of guilt-by-association, where network topology is used to infer gene functions and disease associations^25–27^. The Random Walk with Restart (RWR) algorithm builds on this by simulating a stochastic process that iteratively explores the global structure of the network, repeatedly returning to a set of seed nodes to assess their global influence^28–30^. This method captures both local and global network structure, enhancing predictive power. RWR has been successfully extended to heterogeneous networks, such as disease-gene associations^31^, and to multiplex heterogeneous networks^32,33^, where distinct interaction types are modeled across multiple layers, further improving biological relevance and prediction accuracy.

Building upon this framework, *PDDRNet-MH* represents a conceptual and technical advance by integrating four interlinked multiplex networks representing patients, cell lines, drugs, and genes (**Figure 1**). Each multiplex network is comprised of four **similarity-based layers** that capture distinct biological or pharmacological relationships among entities of the same type. In the patient multiplex, layers encode **patient-patient similarity** based on gene expression, copy number alterations, DNA methylation, and clinical features. The cell line multiplex includes similarity layers based on gene expression, copy number alterations, DNA methylation, and drug screening profiles. The drug multiplex models **drug-drug similarity** using chemical structure, side effects, *in-vitro* drug response, and curated drug interaction data. The gene multiplex represents **gene-gene similarity** through co-expression, protein-protein interactions, and functional similarities derived from Gene Ontology (GO)^34^ annotations. To connect these multiplex networks, five bipartite networks capture inter-entity relationships: patient-cell line similarity (based on cancer-related pathway-level comparisons of multi-omics profiles^35^), cell line-drug sensitivity, curated drug-gene associations, gene-patient mutation profiles, and patient-drug clinical outcomes. This integrated framework enables the propagation of molecular, pharmacological, and clinical signals across the network. By applying the RWR algorithm to this unified architecture, *PDDRNet-MH* predicts drug-patient associations under the assumption that biologically and pharmacologically similar entities are more likely to interact. Detailed descriptions of multiplex layers, bipartite network construction, data processing, and network statistics are provided in the **Materials and Methods** and **Supplementary Methods sections**.

**Figure 1.**
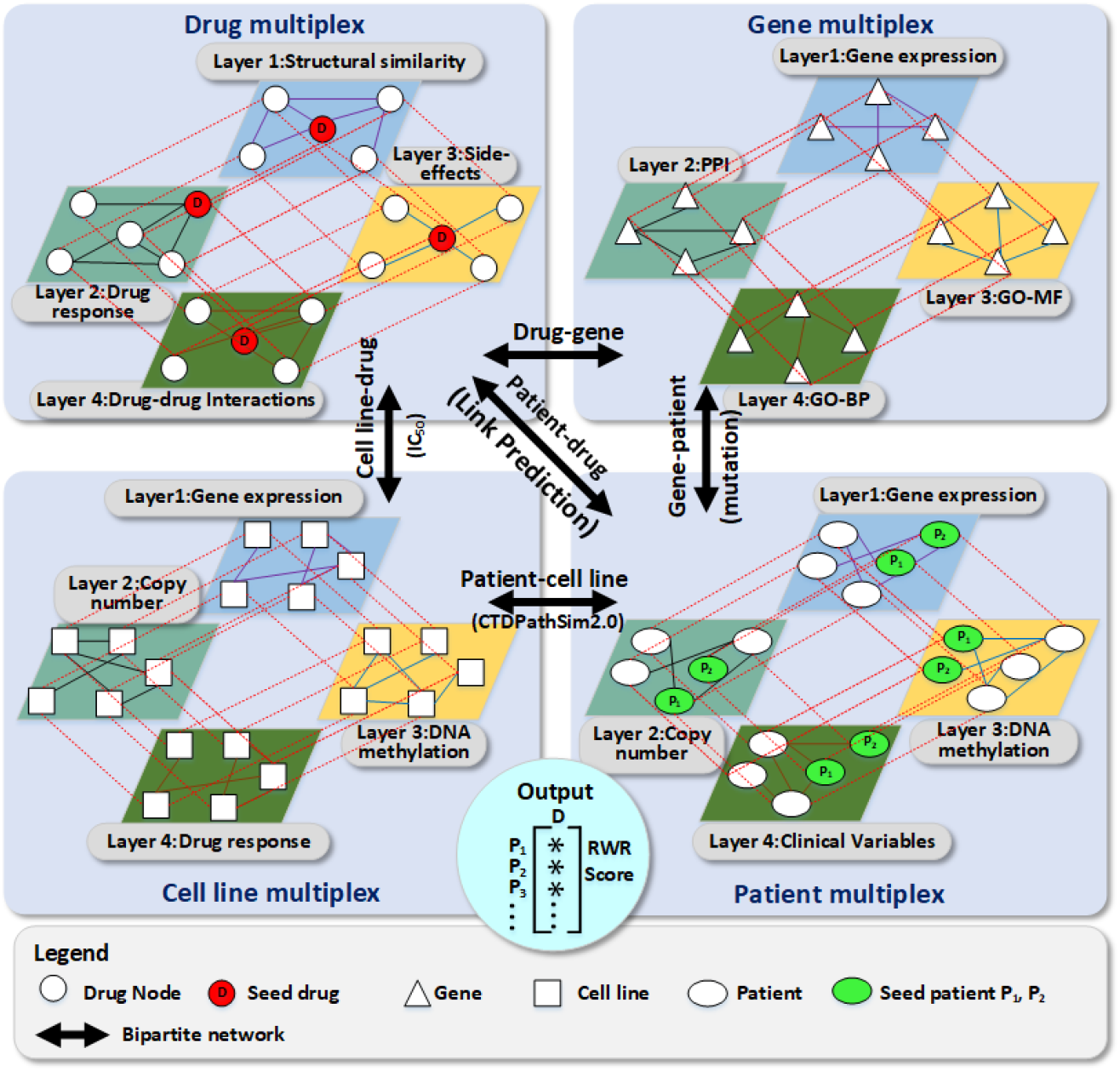
Overview of the *PDDRNet-MH* Framework. *PDDRNet-MH* integrates four multiplex networks-patients, cell lines, drugs, and genes-each comprising four biologically distinct, similarity-based layers. These layers include gene expression, copy number alterations, DNA methylation, clinical variables, structural similarity, drug response, side-effect profile, protein-protein interactions, and gene ontology (GO) annotations. The multiplex networks are interconnected via five bipartite graphs representing patients and cell lines, cell lines and drugs, drugs and genes, genes and patients, and patients and drugs. Red edges indicate intra-multiplex connections within a network, while black edges indicate inter-network bipartite links. Green-colored patient nodes (P1, P2) and red-colored drug nodes (D) serve as seed nodes for initiating the random walk with restart (RWR) process. The final output is a ranked list of drug-patient proximity scores, derived from the steady-state distribution of the RWR process across the multiplex heterogeneous network, enabling personalized drug response prediction through link prediction.

## Results

Application to breast cancer patients in TCGA: We applied *PDDRNet-MH* to predict drug responses in primary breast cancer (BRCA) patients using primary tumor data from The Cancer Genome Atlas (TCGA)^36^. The drug network was built from 397 anticancer compounds profiled across 1,019 cancer cell lines from the Cancer Cell Line Encyclopedia (CCLE)^12^ and the Genomics of Drug Sensitivity in Cancer (GDSC)^11^ projects. All compounds were included in the drug multiplex, but for performance evaluation, we focused on eleven FDA-approved drugs for breast cancer treatment that had sufficient label balance, defined as at least two BRCA patients classified as sensitive and two as resistant. To assess model performance, we performed leave-one-out cross-validation (LOOCV) for each of the eleven drugs (**Materials and Methods**). For each held-out patient, the model generated drug response predictions (score using RWR proximity values (hereafter, RWR scores) between the patient and each drug. These scores were then used to compute receiver operating characteristic (ROC) and precision-recall (PR) curves, evaluating the model’s predictive accuracy.

### *PDDRNet-MH* demonstrates improved predictive power over existing patient-specific drug response models

To assess the predictive performance of *PDDRNet-MH*, we benchmarked it against two established machine learning-based approaches: Imputed Drug-Wide Association Study (IDWAS)^18^ and Support Vector Machine with Recursive Feature Elimination (SVM-RFE)^19^ (**Supplementary Methods**). Unlike many drug response models that rely exclusively on *in vitro* cell line data, both IDWAS and SVM-RFE have been applied to predict drug response in actual patients, making them clinically relevant comparators for our evaluation. IDWAS imputes drug response by leveraging patient transcriptomic profiles, while SVM-RFE is a supervised learning method that incorporates feature selection to classify patients as drug-sensitive or resistant. As both methods represent state-of-the-art strategies for predicting patient-specific drug response, they provide an appropriate benchmark for evaluating the performance of *PDDRNet-MH*.

Model performance was evaluated using the area under the ROC (AUC-ROC), which quantifies the trade-off between the true-positive rate (TPR) and false-positive rate (FPR) across different classification thresholds. In this context, the TPR reflects the proportion of truly sensitive cases correctly identified, while the FPR represents the proportion of resistant cases incorrectly predicted as sensitive. A higher AUC-ROC shows stronger discriminatory power to distinguish between sensitive and resistant patients. Given the class imbalance in the drug response data, where the number of sensitive and resistant cases varied considerably across drugs (**Supplementary Table S2**), we also computed the area under the precision-recall curve (AUC-PR) as a complementary evaluation metric. The AUC-PR evaluates the trade-off between precision (positive predictive value) and recall (sensitivity) and is more informative than AUC-ROC in settings with imbalanced class distributions^37,38^.

### Comparative Performance Across Drugs

Table 1 presents the AUC-ROC and AUC-PR values for *PDDRNet-MH*, IDWAS, and SVM-RFE across the selected drugs. For most drugs, *PDDRNet-MH* outperformed both baseline models, often by a substantial margin. Notably, *it* achieved perfect prediction performance for gemcitabine and vinorelbine (AUC-ROC = 1.00, AUC-PR = 1.00), demonstrating its strong ability to capture meaningful patient-drug associations. For methotrexate and zoledronate, *PDDRNet-MH* also showed near-perfect accuracy (AUC-ROC = 0.95, AUC-PR = 0.99), outperforming IDWAS and SVM-RFE by over 0.4 in AUC-ROC and 0.2 in AUC-PR. On docetaxel, a commonly used chemotherapeutic, *PDDRNet-MH* achieved an AUC-ROC = 0.76 and AUC-PR = 0.97, substantially higher than IDWAS (AUC-ROC = 0.29, AUC-PR = 0.86) and SVM-RFE (AUC-ROC = 0.37, AUC-PR = 0.87). In the case of 5-fluorouracil, IDWAS slightly outperformed *PDDRNet-MH* in AUC-ROC (0.59 vs. 0.58), but both methods had similarly high AUC-PR values (0.91). Epirubicin was the only drug where IDWAS achieved the highest performance (AUC-ROC = 0.74, AUC-PR = 0.98), though *PDDRNet-MH* remained competitive (AUC-ROC = 0.59, AUC-PR = 0.96). For other drugs, including cyclophosphamide, doxorubicin, paclitaxel, and tamoxifen, *PDDRNet-MH* consistently demonstrated superior predictive accuracy across both metrics. The occasional advantages of IDWAS may reflect overfitting to gene expression features, rather than broadly generalizable performance.

Overall, *PDDRNet-MH* outperformed existing methods across the majority of therapeutic agents, demonstrating robust predictive accuracy, higher precision, and improved generalizability to real-world patient data. AUC-ROC and AUC-PR values for eleven breast cancer drugs evaluated on TCGA BRCA samples. **Bolded values indicate the highest performance for each metric per drug**.

**Table 1.**
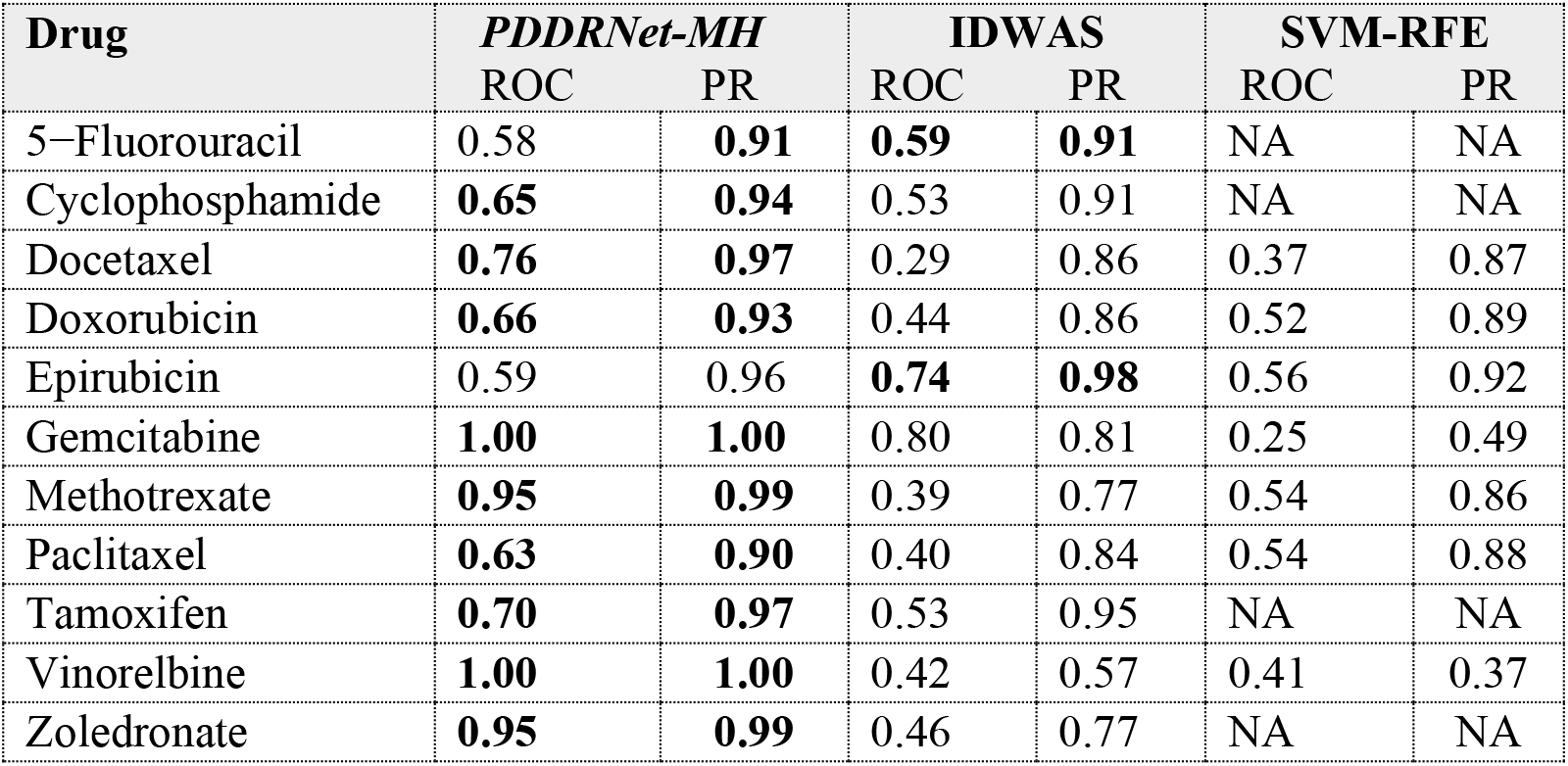
AUC-ROC and AUC-PR values of eleven drugs evaluated on TCGA BRCA samples. Bolded values indicate the highest performance for each metric per drug.

### *PDDRNet-MH* accurately prioritizes HER2-targeted therapy in breast cancer patients

HER2, also known as ERBB2 (*erythroblastic oncogene B2*), is a member of the human epidermal growth factor receptor (EGFR/ERBB) family of receptor tyrosine kinases. Amplification or overexpression of ERBB2 occurs in approximately 15–20% of breast cancers and is associated with more aggressive disease and poorer prognosis^39–41^. Due to its oncogenic potential, HER2 has become a critical therapeutic target. In the TCGA BRCA cohort, samples were categorized as HER2-positive (HER2+), HER2-negative (HER2−), or HER2-equivocal, based on immunohistochemistry scoring (IHC)^42^. The HER2-equivocal includes tumors with indeterminate HER2 status that cannot be definitively classified by IHC alone.

Lapatinib, an oral dual tyrosine kinase inhibitor targeting HER2, is an FDA-approved therapy for HER2+ breast cancers^43–45^. To evaluate whether *PDDRNet-MH* could recover this known clinical association, we performed Leave-one-out Cross-validation (LOOCV) using lapatinib and two HER2+ sensitive patients as seed nodes, following the procedure described in the **Materials and Methods** (with restart parameter *r* = 0.9). We then compared the resulting RWR scores between HER2+ and HER2− groups.

As expected, HER2+ patients exhibited significantly higher RWR scores compared to HER2− patients (Wilcoxon rank-sum p-value = 5.74 × 10−7), indicating that *PDDRNet-MH* effectively prioritizes the known responsive population (**Figure 2**). Notably, HER2-equivocal samples showed intermediate RWR scores between HER2+ and HER2− groups, consistent with their indeterminate clinical classification. Further analysis of the RWR score distribution revealed significant enrichment of HER2+ tumors in the top 10th percentile (hypergeometric p-value = 0.0018), while HER2− tumors were significantly enriched in the bottom 20th percentile (p-value = 7.33 × 10−9). These results demonstrate that *PDDRNet-MH* accurately identifies patients likely to benefit from HER2-targeted therapy.

**Figure 2.**
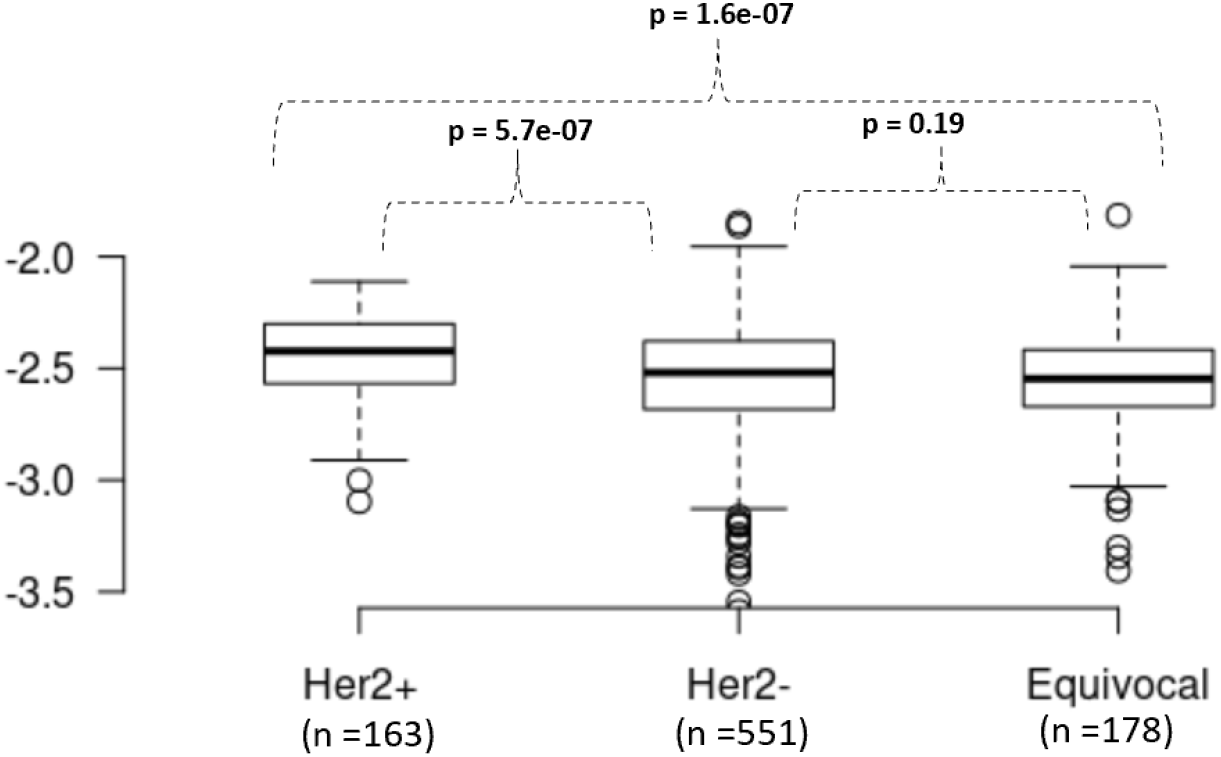
Boxplots of long-transformed RWR scores for lapatinib in TCGA breast cancer samples, stratified by HER2 status as determined by immunohistochemistry. HER2+ tumors exhibited significantly higher scores than HER2− patients (*p* = 5.7 × 10−7), consistent with the known clinical efficacy of lapatinib in HER2+ breast cancer.

To assess the specificity of this association, we examined *PDDRNet-MH* scores for HER2+ and HER2− tumors across the remaining eleven drugs that do not directly target HER2 (**Table 2**). As expected, no significant differences were observed (p-value > 0.05), supporting that the model’s signal for lapatinib reflects a true biological and clinically validated association, rather than an artifact of sample stratification.

**Table 2:**
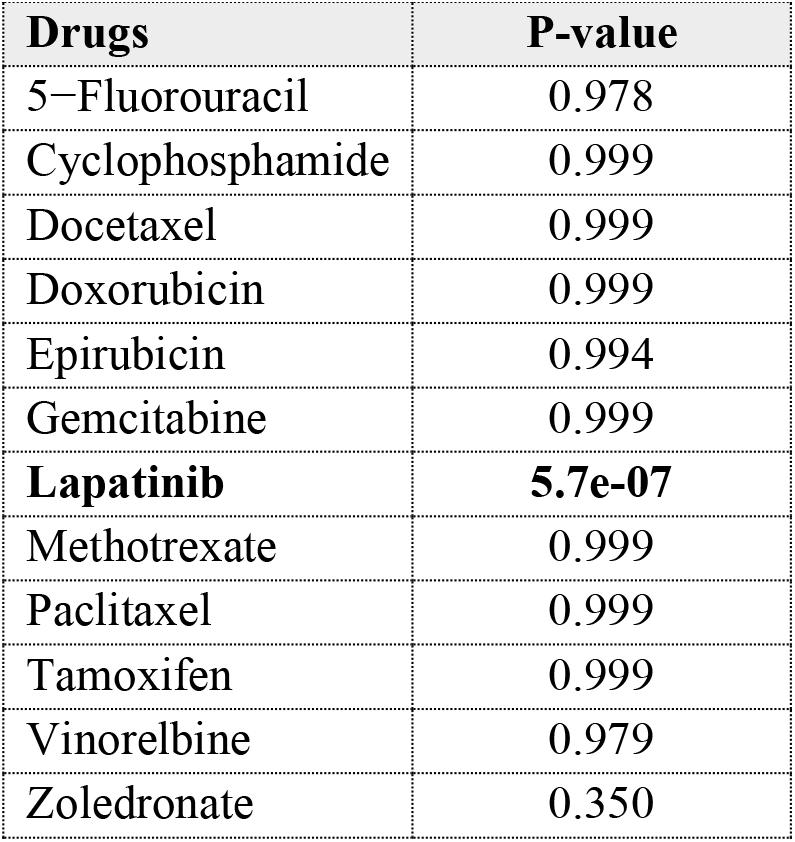
**P-values** for the comparison of RWR scores between HER2+ and HER2− breast cancer tumors in TCGA for each of the **twelve drugs tested** Only lapatinib, a HER2-targeted therapy, showed a statistically significant difference, with higher scores in HER2+ patients, aligning with its established clinical selectivity.

| Drugs | P-value |
| --- | --- |
| 5-Fluorouracil | 0.978 |
| Cyclophosphamide | 0.999 |
| Docetaxel | 0.999 |
| Doxorubicin | 0.999 |
| Epirubicin | 0.994 |
| Gemcitabine | 0.999 |
| **Lapatinib** | **5.7e-07** |
| Methotrexate | 0.999 |
| Paclitaxel | 0.999 |
| Tamoxifen | 0.999 |
| Vinorelbine | 0.979 |
| Zoledronate | 0.350 |

### *PDDRNet-MH* prioritizes established gene biomarkers relevant to drug response

Mutations in the BRCA1 and BRCA2 genes (hereafter, referred to as BRCA1/2) are prevalent in both hereditary and sporadic breast cancers and are associated with defects in DNA repair through homologous recombination. As tumor suppressors, loss of BRCA1/2 function contributes to genomic instability and tumor progression^46,47^. Chemotherapeutic agents such as doxorubicin and cyclophosphamide, which exert their effects through DNA damage, have shown heightened efficacy in BRCA1/2-mutated tumors due to their increased vulnerability to genotoxic stress^48^.

To evaluate whether *PDDRNet-MH* can recover clinically validated gene-drug associations, we examined the RWR scores assigned to BRCA1 and BRCA2 in the gene multiplex network in the context of doxorubicin and cyclophosphamide (**Materials and Methods**). For both drugs, BRCA1 and BRCA2 ranked within the top 3rd percentile of all genes, specifically, among the top 130 genes out of 4,313, indicating strong inferred associations. These results demonstrate that *PDDRNet-MH* effectively recovers established pharmacogenomic markers through its network-based multi-omics integration.

To further assess the model’s ability to identify established drug-gene relationships, we performed a similar analysis using lapatinib, a HER-2 targeted therapy. The ERBB2 gene, which encodes HER2, ranked within the top 1st percentile, placing it among the top 44 genes in the gene multiplex network. These results underscore the ability of *PDDRNet-MH* to accurately prioritize clinically relevant biomarkers, even in a fully data-driven, unsupervised setting.

### *PDDRNet-MH* prioritizes doxorubicin sensitivity in BRCA1/2-mutated triple-negative breast cancer

Breast cancer is clinically and molecularly heterogeneous and is commonly stratified into subtypes based on the expression of estrogen receptor (ER), progesterone receptor (PR), and HER2 (ERBB2). While receptor-positive subtypes often benefit from targeted therapies, triple-negative breast cancer (TNBC), which lacks expression of all three receptors, remains a major therapeutic challenge^49^.

Previous studies have shown that TNBC patients with BRCA1 mutations, and to a lesser extent BRCA2 mutations, may exhibit increased sensitivity to doxorubicin, an anthracycline chemotherapeutic^50,51^. To assess whether *PDDRNet-MH* captures this mutation-specific sensitivity, we analyzed the RWR scores computed for doxorubicin (**Materials and Methods**) in TNBC patients stratified by BRCA mutation status: TNBC with BRCA1 and/or BRCA2 mutations (Group 1) and TNBC patients without BRCA1/2 mutations (Group 2).

Group 1 exhibited significantly higher RWR scores for doxorubicin than Group 2 (p-value = 0.017), indicating that *PDDRNet-MH* reflects known mutation-specific therapeutic sensitivities in TNBC for doxorubicin. This supports the utility of *PDDRNet-MH* in enabling biologically informed patient stratification in the context of personalized therapy selection.

### *PDDRNet-MH* captures drug-associated somatic copy number alterations in the HER2 amplicon

Building on the observed association between lapatinib RWR scores and HER2 status in breast cancer patients, we next investigated whether this signal could be attributed to somatic copy number alterations (CNAs) in the HER2 amplicon. Overexpression of HER2 is frequently driven by copy number amplification within a well-characterized region on chromosome 17q12, commonly referred to as the amplicon^52,53^. If *PDDRNet-MH* could identify this amplification region as a predictive of drug response, it would underscore the model’s utility in uncovering functionally relevant somatic alterations.

To explore this, we analyzed copy number segmentation profiles from TCGA BRCA samples using GISTIC 2.0^54^ in GenePattern^55^. We extracted amplification status for 129 genes located within the 17q12 HER2 amplicon using a confidence threshold of 0.90, categorizing samples into CNA-amplified and non-amplified groups for each gene.

Using lapatinib and two sensitive patients as seeds in the network, we applied the RWR algorithm to compute gene-level RWR scores (**Materials and Methods**). For each of the 129 genes in the HER2 amplicon, we compared RWR scores between CNA-amplified and non-amplified samples. Strikingly, 101 out of 129 genes showed significantly higher RWR scores in the amplified group (*Wilcoxon rank-sum p-value < 0*.*05*), indicating that *PDDRNet-MH* captures consistent CNA-associated drug relevance in this region. Notably, HER2 itself exhibited a significantly elevated RWR score in the amplified group (p-value = 0.003; **Figure 3**), validating the model’s alignment with known drug-gene relationships.

**Figure 3.**
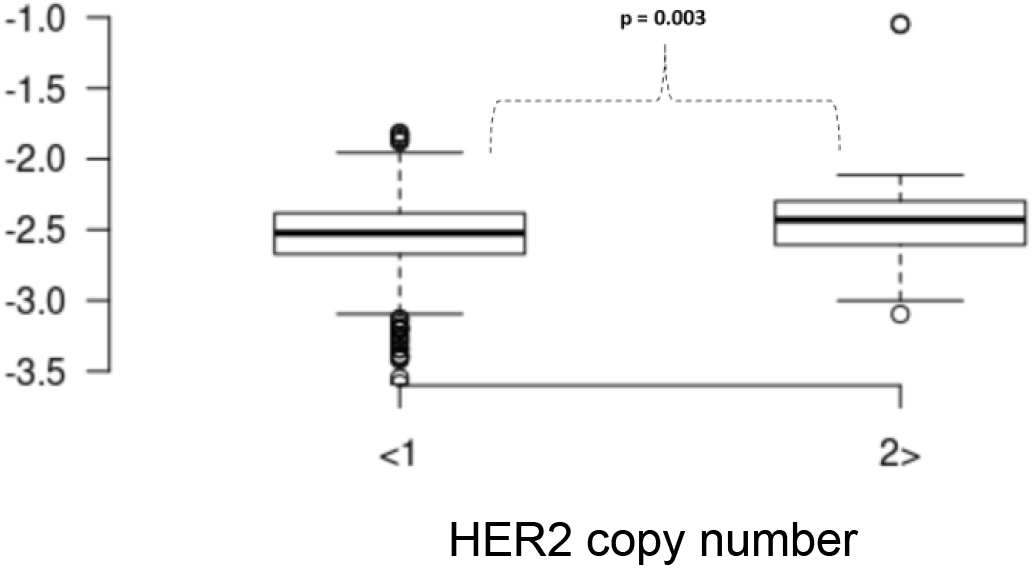
Boxplots of log-transformed RWR scores for genes in TCGA breast cancer samples, stratified by HER2 copy number status. Samples with HER2 amplification exhibited significantly higher RWR scores compared to non-amplified samples (*p*-value = 0.003).

To prioritize candidate biomarkers within the HER2 amplicon, we selected the top 10th percentile of genes based on both effect size and statistical significance (**Table 3**). Remarkably, HER2 ranked among the top six genes, along with PPP1R1B, STARD3, and MIEN1, genes previously implicated in HER2 signaling and lapatinib sensitivity in breast cancer^56–59^. These findings illustrate that *PDDRNet-MH* can identify multiple functionally relevant genes within amplified regions, providing a data-driven pathway for discovering novel genomic predictors of drug response.

**Table 3.** Top-ranked genes within the HER2 amplicon for which the RWR scores for lapatinib significantly differ between CNA-amplified and non-amplified patient groups in TCGA BRCA. Genes are ranked by p-value and effect size. Literature references support their known or potential relevance to HER2^+^ breast cancer or lapatinib response.

| Gene | No. Of. amplified samples | No. Of. not-amplified samples | P-value | Effect-size | Supporting literature |
| --- | --- | --- | --- | --- | --- |
| NEUROD2 | 117 | 774 | 2.1e-04 | 0.259 | Lacle et al. 2015 |
| PPP1R1B | 121 | 769 | 3.8e-05 | 0.250 | Ma et al. 2018; Sahlberg et al. 2013 |
| STARD3 | 122 | 766 | 4.4e-05 | 0.247 | McDermott et al. 2013, 2; Ma et al. 2018 |
| TCAP | 122 | 766 | 4.4e-05 | 0.247 | Serie et al. 2017 |
| PNMT | 122 | 766 | 4.4e-05 | 0.247 | Arriola et al. 2008; Khoury et al. 2010, 2 |
| PGAP3 | 124 | 765 | 4.4e-05 | 0.243 | Ferrari et al. 2016 |
| HER2 | 124 | 765 | 4.4e-05 | 0.243 | Ramlau et al. 2015; Johnston and Leary 2006; F. Yang et al. 2020 |
| MIEN1 | 121 | 769 | 9.4e-05 | 0.194 | Rexer and Arteaga 2012, 2 |
| GRB7 | 121 | 768 | 9.4e-05 | 0.194 | Rexer and Arteaga 2012, 2 |
| IKZF3 | 118 | 772 | 1.4e-04 | 0.200 | Sircoulomb et al. 2010, 2-; Ye et al. 2019 |

## Discussion

We developed *PDDRNet-MH*, a computational framework for predicting patient-specific drug responses by integrating multi-omics and pharmacological data within a multiplex heterogeneous network model. This framework incorporates four multiplex networks: patients, cell lines, drugs, and genes, each comprised of multiple layers representing distinct biological or chemical similarity measures. These networks are interconnected through five types of bipartite edges, enabling the propagation of information across biologically meaningful relationships.

PDDRNet-MH employs the **Random Walk with Restart (**RWR) algorithm, which simulates a stochastic walker starting from a set of seed nodes. The walker moves across intra- and inter-network layers, guided by global and layer-specific transition probabilities, and periodically restarts at seed nodes with a defined probability. This approach allows *PDDRNet-MH* to capture both local and global topological structures, making it particularly well suited for integrating diverse biological signals into a unified predictive framework.

When applied to breast cancer data, *PDDRNet-MH* successfully recovered known associations between targeted therapies and molecular subtypes. It assigned higher drug relevance scores to patients with established biomarkers such as HER2 amplification and BRCA1/2 mutations. Moreover, it was able to accurately identify mutation-specific drug sensitivity in difficult-to-treat subtypes like triple-negative breast cancer. It also captured drug-gene associations consistent with clinical knowledge.

Importantly, beyond capturing the effects of established biomarkers, *PDDRNet-MH* also prioritized genes within recurrent somatic copy number alterations that correlate with drug response. For example, it highlighted genes co-amplified with HER2 in chromosome 17q12 that are implicated in resistance or sensitivity to HER2-targeted therapies. These results suggest that *PDDRNet-MH* not only validates established pharmacogenomic relationships but also has the potential to uncover novel genomic alterations with therapeutic relevance, supporting its utility in precision oncology.

To assess the contribution of individual data components, we conducted ablation experiments by selectively removing specific multiplex networks. Excluding the cell line network led to a noticeable decline in predictive performance across most drugs, indicating its critical role in linking pharmacological properties to patient molecular profiles. The gene network also contributed to model accuracy, albeit to a lesser degree. These results underscore the importance of incorporating multi-modal biological knowledge into the network to capture the multifactorial nature of drug response in patients.

We also evaluated the model’s sensitivity to key RWR parameters. Adjusting the global restart probability revealed improved performance when the random walker was increasingly constrained to the local neighborhood of the seed nodes, suggesting that localized signal propagation enhances predictive specificity. Additionally, varying the inter-layer transitions within multiplex networks, as well as the node-level restart probabilities for the patient and drug networks, impacted model performance. The patient multiplex network emerged as particularly influential; its layered integration of gene expression, somatic mutations, DNA methylation, and clinical similarity contributed significantly to prediction accuracy. Together, these findings emphasize the importance of balanced, integrative network design and careful parameter tuning when modeling complex phenotypes such as drug response in a biologically meaningful way.

Although *PDDRNet-MH* demonstrated strong predictive performance and robustness, several limitations remain. The current evaluation was limited to a subset of drugs with sufficient numbers of sensitive and resistant patients to enable leave-one-out cross-validation. However, this constraint stems from the validation strategy, not from the model itself, which is fully capable of prioritizing drug responses in unlabeled or less well-characterized patient cohorts. In addition, while *PDDRNet-MH* integrates diverse multi-omics data layers, it does not yet account for tumor microenvironmental features such as immune infiltration, stromal interactions, or spatial heterogeneity, factors increasingly recognized as key modulators of treatment response. Future extensions of *PDDRNet-MH* will incorporate such context-specific factors and explore alternative seeding strategies, including the use of drug or gene nodes alone, to enhance model generalizability and clinical utility.

In summary, *PDDRNet-MH* offers a biologically grounded, computationally robust framework for predicting therapeutic responses in cancer. Integrating multi-omics data within a network-based learning model enables scalable and interpretable biomarker discovery and personalized drug prioritization. Broader application across multiple cancer types and integration with clinical outcomes may further enhance its utility in translational precision oncology.

## Materials and Methods

### Multiplex and bipartite network construction

We developed a multiplex heterogeneous network framework to predict patient-specific drug responses by integrating four types of biological and pharmacological entities: patients, cell lines, drugs, and genes. Each entity was modeled as a multiplex network consisting of four distinct data-driven similarity layers, and the resulting subnetworks were interconnected by five bipartite networks that capture inter-entity relationships. This integrated structure enables *PDDRNet-MH* to incorporate complementary information across molecular, pharmacological, and clinical domains. While all four multiplex networks were implemented, only the patient and drug networks are essential for the execution of the algorithm; the inclusion of the gene and cell line networks further enhances prediction performance and biological interpretability.

The patient multiplex network models inter-patient similarity using four data modalities: (1) gene expression, (2) CNA, (3) somatic mutations, and (4) clinical phenotypes. The clinical layer integrates both ordinal and nominal clinical variables into a composite similarity matrix, with variable-specific weights derived from Cox regression modeling. Details on variable encoding and data harmonization are provided in **Supplementary Methods** and summarized in **Supplementary Table S1**.

The cell line multiplex network models similarity among cancer cell lines using (1) gene expression, (2) CNA, (3) somatic mutation profiles, and (4) drug response. The drug response layer models pharmacological similarity using normalized sensitivity profiles across a shared drug panel. Data integration procedures and filtering criteria are detailed in **Supplementary Methods**.

The drug multiplex network connects drugs based on (1) chemical structure similarity, (2) drug response similarity in cancer cell lines, (3) side-effect profile similarity, and (4) curated drug-drug interactions. These layers capture the structural, mechanistic, and functional properties of drugs. The derivation of these features from curated databases and molecular descriptors is described in **Supplementary Methods**.

The gene multiplex network represents relationships among cancer-relevant genes and the most variable genes in the BRCA patient transcriptome. It includes layers reflecting (1) gene co-expression patterns, (2) protein-protein interactions, (3) GO-biological process similarity, and (4) GO-molecular function-based similarities. Details on network construction, edge weighting, and gene selection are provided in **Supplementary Methods**. To interconnect these multiplex networks, we constructed five bipartite networks: patient-cell line similarity (based on pathway activity-level comparisons of multi-omics profiles); cell line-drug relationships (reflecting sensitivity or resistance via binarized drug response values); drug-gene associations (derived from curated pharmacogenomic databases); gene-patient mutation profiles (using binary indicators of somatic mutation status); and patient-drug outcomes (defined from clinical response annotations and survival data). Complete procedures for bipartite graph construction, including database harmonization, threshold selection, and edge definition, are described in the **Supplementary Methods**. In each multiplex network, the final set of nodes was defined as the intersection of nodes present in all four layers, ensuring consistent entity representation across modalities. **Supplementary Table S3** summarizes the final numbers of nodes and edges in each multiplex and bipartite network for the breast cancer application of *PDDRNet-MH*.

### Random Walk with Restart (RWR) Computation in *PDDRNet-MH*

*PDDRNet-MH* utilizes the steady-state distribution of RWR to compute the proximity score of the nodes in the network from the seed nodes by using Eq. (1),

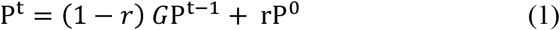

where *r* is the global restart probability (i.e., the likelihood of the surfer returning to the seed nodes at each step), P^t^ is the probability distribution vector of all the nodes at time *t*, P^0^ denotes the probability distribution vector of all the nodes at time *t* = 0, i.e., the restart vector and *G* is the global transition matrix of the multiplex heterogeneous network (see **Supplementary Methods for its construction**) of *PDDRNet-MH*. Equation (1) is solved iteratively by updating P^t^ until convergence, i.e., |P^*i*^ − P^*i*−1^|< ε where *i* denotes the iteration number and ε = 10^−10^ (a standard threshold for the RWR algorithm)^31,32^. P^*i*^ converges to a *unique* solution (P^*s*^), i.e., reaches a steady-state^66^. The steady state distribution (P^*s*^) is used as a proximity vector for the nodes in the network starting from a seed node. Details of the computation of adjacency matrices and random walk in the multiplex heterogeneous network are explained in **Supplementary Methods**.

In addition to the global restart probability (*r*) and inter-layer transition probabilities (*δ*), *PDDRNet-MH* includes eight additional hyperparameters: *τ*_*p*_, *τ*_*c*_, *τ*_*d*_, *τ*_*g*_, *η*_*p*_, *η*_*c*_, *η*_*d*_ and *η*_*g*_. Here *τ*_*p*_, *τ*_*c*_, *τ*_*d*_ and *τ*_*g*_ are the restart probabilities for the multiplex networks, such as patient network (***M***_***PP***_), cell line network (***M***_***CC***_), drug network (***M***_***DD***_), and gene network (***M***_***GG***_), respectively. The default values for these parameters are: 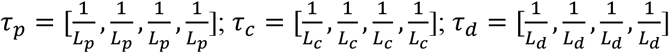 and 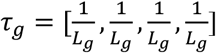; where *L*_*p*_, *L*_*c*_, *L*_*d*_ and *L*_*g*_ denote the number of layers in patient, cell line, drug, and gene multiplex networks, respectively. In this study, we have used *L*_*p*_= *L*_*c*_= *L*_*d*_= *L*_*g*_= 4. The parameters *η*_*p*_, *η*_*c*_, *η*_*d*_ and *η*_*g*_ control the probability of restarting the walk from the seed nodes in each respective network. These weights satisfy the normalization constraint: *η*_*p*_+ *η*_*c*_+ *η*_*d*_+ *η*_*g*_ = 1. To ensure balanced information propagation at initialization, *PDDRNet-MH* sets the layer scaling factor parameter λ = 1/N, where N is the number of multiplex networks, to evenly allocate the initial probability across all networks. This encourages the random walker to form any network with equal likelihood, supporting balanced information propagation across the heterogeneous structure.

### Leave-one-out cross-validation in drug response prediction

We applied *PDDRNet-MH* to predict links between the patient multiplex network (***M***_***PP***_) and the drug multiplex network (***M***_***DD***_) using a leave-one-out cross-validation (LOOCV) strategy. For a given drug D in ***M***_***DD***_, the objective was to evaluate the model’s ability to predict patient-specific sensitivity to D. To do this, we identified all of the patients from ***M***_***PP***_ who were known to be sensitive to drug D. Then, for each such patient P, we withheld the known patient-drug association by removing the corresponding edge between P and drug D in ***B***_***PD***_, setting the corresponding bipartite edge weight to 0. This edge removal ensured that the association was excluded during inference, allowing us to evaluate whether *PDDRNet-MH* could successfully **recover the hidden link** through network-based propagation. Next, we performed RWR on the multiplex heterogeneous network, initializing the random walk with drug D and all remaining patients known to be sensitive to D (excluding the held-out patient P) as seed nodes. The RWR algorithm iteratively propagated proximity scores across the full network using Equation (1), until convergence to a steady-state distribution. We then z-normalized the resulting node scores to account for variation in score distributions and retrieved the RWR score of the held-out patient P. This procedure was repeated for every known sensitive patient-drug pair, sequentially removing the corresponding edge in ***B***_***PD***_ and retrieving the respective score of the held-out patient. This iterative leave-one-out removal and prediction defines our LOOCV strategy. Separately, we also retained z-normalized scores for unknown and resistant patients, obtained from an RWR run using all known sensitive patients for drug D as seed nodes without removing any edges. Z-normalization helps mitigate score distribution shifts that might arise due to changes in the seed set, allowing for fair comparison between sensitive, resistant, and unlabeled patients.

This cross-validation procedure was performed separately for each drug. For these experiments, we used the following default hyperparameter values^33^: global restart probability *r* = 0.7; inter-layer walk probability *δ* = 0.5; layer scaling factor *λ* = 0.25; and multiplex restart weights *η*_*p*_ = 0.75, *η*_*d*_ = 0.25, with *η*_*d*_ = 0; *η*_*g*_ = 0, since no seed nodes were selected from the cell line or gene networks during these tests.

To evaluate the robustness and contribution of individual components, we performed a series of ablation studies and parameter sensitivity analyses. These included comparisons of LOOCV performance with and without the cell line and gene networks, as well as selective removal of individual layers within each multiplex network. Removing either the gene or cell line network led to a notable drop in predictive accuracy, confirming their complementary roles in capturing biologically meaningful information. Among the individual layers, the clinical and transcriptomic similarity layers in the patient network and the structural similarity layer in the drug network contributed most significantly to model performance. To assess model robustness, we systematically varied key hyperparameters, including the global restart probability, multiplex-specific restart probability vectors, and inter-multiplex restart weights, and found that the model maintained stable prediction accuracy across a wide range of values, highlighting its’ resilience to parameter variations. Full details and results of these evaluations, including results and implementation details, are provided in **Supplementary Methods**.

### Computing the RWR scores for the gene nodes associated with a drug

To evaluate whether *PDDRNet-MH* could recover established gene-drug associations, we computed RWR scores for BRCA1 and BRCA2 in the gene multiplex network using doxorubicin and its associated sensitive patients as seed nodes. We repeated the analysis separately using cyclophosphamide and its corresponding sensitive patients as seed nodes. To prevent the risk of direct or indirect bias, we removed all bipartite edges connecting any gene node to either of the seed drug or its associated sensitive patients in the drug-gene and gene-patient bipartite networks. This ensured that gene prioritization was not influenced by pre-existing associations with the seed nodes. We then applied the RWR algorithm with a restart probability of *r* = 0.9, using the drug and its known sensitive patients as seed nodes, to compute steady-state proximity scores for all genes in the network. These RWR scores reflect the inferred network proximity of each gene to the seeded drug and patient nodes, independent of any direct connections. The resulting ranked gene lists were used to evaluate whether *PDDRNet-MH* could correctly prioritize BRCA1 and BRCA2 as known pharmacogenomic markers. A similar procedure was applied to compute RWR scores for genes within the HER2 amplicon, using lapatinib and its associated sensitive patients as seed nodes.

## Data and Code Availability

The full source code for *PDDRNet-MH*, including scripts for network construction, Random Walk with Restart, and model evaluation, is available at: https://github.com/bbose/PDDRNet_MH. Breast cancer molecular and clinical data were obtained from TCGA via the Genomic Data Commons (GDC) Data Portal: https://portal.gdc.cancer.gov/. Cancer cell line drug response data and molecular features were obtained from CCLE: https://portals.broadinstitute.org/ccle and GDSC: https://www.cancerrxgene.org/. Data processing details are provided in the Supplementary Methods.

## Acknowledgments

This work was supported by the National Institute of General Medical Sciences of the National Institutes of Health under Award Number R35GM13365, and by the National Cancer Institute of the National Institutes of Health under Award Number R01CA229618.

