## Supplementary Methods for "A network model for patient-derived drug response in breast cancer integrating multi-omics datasets"

In this work, we constructed a multiplex heterogeneous network model that consists of patient, cell line, drug, and gene multiplex networks (**Figure 1 in the main manuscript**) to predict patient-specific drug response. In each multiplex network, the final set of nodes was defined as the intersection of nodes present across all four constituent layers to ensure consistent entity representation. Five bipartite networks were considered to construct the edges between these networks: patient-cell line, cell line-drug, drug-gene, gene-patient, and patient-drug. We introduced the construction of these multiplex and the bipartite networks in the following sections. Although we utilized the four multiplex networks in *PDDRNet-MH*, this tool could be run without cell line and gene networks.

**Building patient multiplex network**

The patient multiplex network ($\boldsymbol{M_{PP}}$) was constructed with four different layers of patient-patient similarity networks. $\boldsymbol{M_{PP}}$ can be considered as an undirected graph, $M_{PP} = (V_P, E_P)$, where $V_P$ denotes a set of patients as vertices (i.e., nodes), and $E_P$ denotes a set of paired patient vertices (i.e., edges) from all the layers. Following is the construction of each layer.

***Layer 1:*** **Gene expression.** This layer was built with patient-patient similarity edges constructed from gene expression profiles. We downloaded mRNA data in Fragments Per Kilobase of transcript per Million mapped reads (FPKM) for 1,064 breast invasive carcinoma (BRCA) tumor samples from the TCGA database. Since FPKM values were skewed, we converted this data matrix using log2(FPKM+1) and ranked all the genes according to their expression variation across all the samples. The 1,000 most variable genes were kept as "marker genes" to compute the absolute Spearman rank correlation ($S_{rc}$) scores as edge weights. We tried different gene sizes in the early preliminary analysis and did not observe a large variation of results, so we decided to choose the 1,000 most-varied genes. This approach is common in gene expression-based studies[1,2].

***Layer 2:*** **Copy number.** This layer was built with patient-patient similarity edges constructed from CNA profiles. We downloaded the CNA segmented data in Affymetrix SNP Array 6.0 for 1,064 BRCA samples from the TCGA database. We have utilized the genomic location information of 22,310 protein-coding genes provided by GENCODE Release 26[3] and applied the R Bioconductor package *CNTools*[4] to convert this segmented CNA data into gene-centric copy number values. We ranked all the genes according to their gene-centric copy number variation across all the tumor samples. The 1,000 most variable genes were kept for computing $S_{rc}$ scores as edge weights.

***Layer 3:*** **DNA methylation.** This layer was built with patient-patient similarity edges constructed from DNA methylation profiles of 1,064 BRCA samples. We downloaded Infinium HumanMethylation27 Bead-Chip (27K) and Infinium HumanMethylation450 Bead-Chip (450K) platform-based DNA methylation data from the TCGA database. Gene-specific beta values were calculated separately for both platforms. For the 450K platform, the average beta value for promoter-specific probes was considered due to their role in transcriptional silencing[5]. Given lower coverage in the 27K platform, we utilized all the probes. In this case, we set the DNA methylation of a gene as the average of the beta values of all its probes. We ranked all the genes according to their beta value variation across all the samples. The 1,000 most variable genes were kept for computing $S_{rc}$ scores as edge weights.

***Layer 4:*** **Clinical variables.** This layer was built with patient-patient similarity edges constructed from 1,064 BRCA patients' clinical data from TCGA. We downloaded the clinical data with two ordinal variables, 'age' and 'stage' (i.e., tumor stage), and eight nominal variables, such as 'race', 'HER2', 'estrogen', 'progesterone', 'Overall Survival (OS)', 'Progression Free Interval (PFI)', 'Disease-Specific Survival (DFI)' and 'Disease Free Interval (DSS)'. In OS, patients who were dead from any cause were considered dead, otherwise censored. In PFI, patients having new tumor event whether it was a progression of the disease, local recurrence, distant metastasis, new primary tumor event, or died with cancer without a new tumor event, including cases with a new tumor event whose type is N/A were considered as "event occurred" and all other patients were censored. DFI was similar to PFI with the inclusion of censored patients with new primary tumors in other organs; patients who were dead with tumors without new tumor event and patients with stage IV were excluded. In DSS, disease-specific survival time in days, last contact days, or death days, whichever was larger, was used to identify "event occurred" versus censored patients[6]. We had four categories for variable 'age' and nineteen different categories for the variable 'stage'. We had five classes for the variable 'race'. **Supplementary Table S1** shows the different clinical variables with the corresponding categories and classes.

**Supplementary Table S1: Clinical variables of TCGA BRCA samples used for building the clinical layers of the patient multiplex network.** { } shows the numerical values of the ordinal variables used in the study.

| Type | Clinical variable | Category or class |
|---|---|---|
| **Ordinal** | Age | Old (age ≥ 56); {4}<br>Middle-aged (36 ≤ age < 56); {3}<br>Adult (18 ≤ age < 36); {2}<br>Young (age < 18); {1} |
| | Stage | Stage 0, I/II NOS, IS, Stage I, Stage IA, Stage IB, Stage II, Stage IIA, Stage IIB, Stage IIC, Stage III, Stage IIIA, Stage IIIB, Stage IIIC, Stage IV, Stage IVA, Stage IVB, Stage IVC, Stage X; {1-19, respectively} |
| **Nominal** | Race | White<br>Black or African American<br>Asian<br>American Indian or Alaska Native<br>Native Hawaiian or other Pacific Islander |
| | HER2 | Positive/Negative/Equivocal/Indeterminate |
| | Estrogen | Positive/Negative/Indeterminate |
| | Progesterone | Positive/Negative/Indeterminate |
| | OS | 1/0 (dead/censored) |
| | PFI | 1/0 (false case/true case) |
| | DFI | 1/0 (false case/true case) |
| | DSS | 1/0 (false case/true case) |

We created the patient-patient clinical profile-based similarity layer by computing similarity scores between each pair of patient nodes in three steps. In the first step, we ran a multivariable Cox regression[7] analysis using R package *survival*[8] for OS as survival status and all other clinical variables as covariates. Using the Cox regression z-score for each variable, we computed weight for each clinical variable, i.e., cox-weight $w_c$ such that $\sum_{c=1}^{n} w_c = 1$; where n denotes the total number of clinical variables.

In the second step, we used different similarity measures appropriate to the variable type. Suppose a clinical variable is denoted by a set **C** = $\{c_i\}$, where an element $c_i$ denotes a clinical value (nominal or ordinal) corresponding to patient *i*. For a nominal clinical variable *c*, the similarity score between patient *i* and patient *j* was computed as: $S_{ij}^{c} = 1$, if the patients had the same class label and $S_{ij}^{c} = 0$, if the patients had different class labels. For an ordinal clinical variable *c*, the similarity score between patient *i* and patient *j* was computed as: $S_{ij}^{c} = 1 - \frac{c_i - c_j}{\max(\mathbf{C}) - \min(\mathbf{C})}$. In the final step, we computed an edge weight between patient *i* and patient *j* as: $S_{ij}^{F} = \frac{\sum_{c=1}^{n} w_c \times S_{ij}^{c}}{n}$.

**Building cell line multiplex network**

The cell line multiplex network ($\boldsymbol{M_{CC}}$) was built with four different layers of cell line- cell line similarity networks. $\boldsymbol{M_{CC}}$ can be considered as an undirected graph, $M_{CC} = (V_c, E_c)$, where $V_c$ denotes a set consisting of cell lines as vertices, and $E_c$ denotes a set of paired cell line vertices from all the layers. Following is the construction of each layer.

***Layer 1:*** **Gene expression.** This layer was built with cell line-cell line similarity edges constructed from gene expression profiles. We downloaded the CCLE_RNAseq_gene_rpkm _10180929.gct.gz file from the CCLE database as gene expression quantification for 1,019 cell lines. We ranked all the genes based on their expression variation and selected the 1,000 most variable genes for computing $S_{rc}$ scores as edge weights.

***Layer 2:*** **Copy number.** This layer was built with cell line-cell line similarity edges constructed from CNA profiles. We downloaded gene-centric CNA (Affymetrix SNP Array 6.0) for 916 cell lines from CCLE. We ranked the genes according to their gene-centric copy number values across all the cell lines. The 1,000 most variable genes were kept for computing $S_{rc}$ scores as edge weights.

***Layer 3:*** **DNA methylation.** This layer was built with cell line-cell line similarity edges constructed from the DNA methylation profile. We downloaded the CCLE_RRBS_cgi_CpG_clusters_20181119.txt.gz file for DNA methylation data of 831 cell lines from the CCLE database. Using the R package *BioMethyl*[9], we removed CpG sites that had missing values in more than half of the samples and imputed the rest of the missing values using the k-nearest neighbor approach utilizing the *Enmix*[10] R package with default parameters. We ranked all the genes according to their beta-value variation across all the cell lines. The 1,000 most variable genes were kept for computing absolute $S_{rc}$ scores as edge weights.

***Layer 4:*** **Drug response.** This layer was built with cell line-cell line similarity edges constructed from the drug response profile. We retrieved 936 cell lines from CCLE and GDSC databases screened with 397 drugs. We considered 77 drugs that were tested on at least 95 percent of total cell lines and normalized their $IC_{50}$ values across all the cell lines between 0 to 1. Using these normalized $IC_{50}$ values, we computed the $S_{rc}$ scores as edge weights.

**Building a drug multiplex network**

The drug multiplex network ($\boldsymbol{M_{DD}}$) was built with four different layers of drug-drug similarity networks. $\boldsymbol{M_{DD}}$ can be considered as an undirected graph, $M_{DD} = (V_d, E_d)$, where $V_d$ denotes a set consisting of drugs as vertices, and $E_d$ denotes a set of paired drug vertices from all the layers. Following is the construction of each layer.

***Layer 1:*** **Structural similarity.** This layer was built with drug-drug similarity edges constructed from structural similarities of 376 drugs from the CCLE and GDSC databases. We retrieved the PaDEL descriptor[11] of the drugs' chemical properties and 2D and 3D structural profiles of the drugs using a line notation of a simplified molecular-input line-entry system (SMILES)[12] mined from the PubChem[13] database. We then applied SVD on a z-normalized matrix of PaDEL descriptors. We performed singular value decomposition (SVD) on the z-normalized PaDEL descriptor matrix, treating each row as a drug and each column as a molecular descriptor. We retained the first five left singular vectors (eigenarrays), which capture the principal components of structural variability among drugs. The resulting matrix provided a low-dimensional representation of each drug, and the absolute Pearson correlation ($P_c$) between these vectors was used as the structural similarity score between drugs.

***Layer 2:*** **Drug response.** This layer was built with drug-drug similarity edges constructed from cell line-specific drug response values from CCLE and GDSC. We considered 244 cell lines that were screened by at least 95 percent of the total 397 drugs. We performed the z-normalization on the cell line-specific $IC_{50}$ values (the concentration of a drug at which

50% of its maximal inhibitory effect is observed) across all chosen drugs. Using these $IC_{50}$ values, we computed $S_{rc}$ scores as edge weights.

***Layer 3:*** **Side-effects.** This layer was built with drug-drug similarity edges constructed from drug side-effects data. We downloaded the drug side effects from the SIDER[14] and OFFSIDES[15] databases. SIDER contains information on adverse drug reactions for FDA-approved drugs. OFFSIDES contains information on side effects that are not listed on the FDA's official drug labels. We retrieved 1,580 side-effect terms from SIDER and OFFSIDES for 57 drugs that were common in the drugs retrieved from CCLE and GDSC. For two drugs $i$ and $j$, the Jaccard score was calculated as $J_s = \frac{|SE_i \cap SE_j|}{|SE_i \cup SE_j|}$, where $SE_i$ and $SE_j$ denote side-effect sets that were belong to $i$ and $j$, respectively. The Jaccard score between each pair of drug nodes was considered as edge weight.

***Layer 4:*** **Drug-drug interaction.** This layer was built with drug-drug interaction data. We retrieved drug-drug interaction data from the Drugbank[16] database for 33 drugs with 910 interaction edges. In Drugbank, an interaction edge between two drugs was considered when they interact, interfere, or cause adverse reactions when taken with each other. We used an edge weight to be 1 if the two drugs had interaction; otherwise, the edge weight was set to 0.

**Building a gene multiplex network**

The gene multiplex network ($\boldsymbol{M_{GG}}$) was built with four different layers of gene-gene similarity networks. $\boldsymbol{M_{GG}}$ can be considered as an undirected graph, $M_{GG} = (V_g, E_g)$, where $V_g$ denotes a set of genes as vertices, and $E_g$ denotes a set of paired gene vertices from all the layers. This network was constructed using 3,313 cancer-related genes and 1,000 most variable genes chosen from the patient samples. We considered cancer-related and patient-specific most varied genes to allow their effect on drug response prediction. The cancer genes were mined from Cancer Gene Census in the COSMIC database[17], and Network of Cancer Genes 5.0[18]. Following is the construction of each layer.

***Layer 1:*** **Gene expression.** This layer was built with gene-gene similarity edges constructed from patient-specific gene expression profiles (*Layer 1*) by computing $S_{rc}$ scores as edge weights.

***Layer 2:*** **PPI.** This layer was built with gene-gene similarity edges constructed from protein-protein interaction (PPI) data downloaded from the STRING database[19], which hosts both experimentally validated and computationally predicted PPIs. Here, species limited to "Homo sapiens" with interaction scores > 0.3 were applied to construct the PPI networks. This threshold was chosen to select confident interactions among the proteins. We further normalized the PPI scores ($S_{PPI}$) between 0 to 1 as edge weights using minmax normalization.

***Layer 3*** and ***Layer 4*** of gene multiplex were constructed with gene ontology (GO) terms[20]. GO provides ontologies to annotate gene products. We utilized the R package *GoSemSim*[21] to compute semantic similarities between two GO terms. In particular, two GO terms were considered to be similar if they were close to each other in the GO-directed acyclic graph (DAG), and two GO terms that were close to each other at a lower level would be considered more similar than those at a higher level in the GO DAG. Several studies suggest that GO terms are informative for characterizing biological mechanisms underlying disease and can provide valuable context for understanding therapeutic responses[22,23]. In this study, we utilized molecular function-related GO (GO-MF) terms and biological processes-related GO (GO-BP) terms. We incorporated two GO-based layers: one using molecular function (GO-MF) terms, which represent molecular-level gene activities such as catalysis or transport[24], and another using biological process (GO-BP) terms, which describe higher-order biological events such as DNA repair or signal transduction[20]. Semantic similarity scores derived from each ontology were used as edge weights in their respective layers.

**Building bipartite networks**

In *PDDRNet-MH*, we constructed five different bipartite relations or bipartite networks to connect the multiplex networks. Following is the construction of each of these bipartite networks:

**Patient-cell line bipartite network.** The patient-cell line bipartite network ($\boldsymbol{B_{PC}}$) was built with patient-cell line similarity networks. $\boldsymbol{B_{PC}}$ can be defined as an undirected graph, $B_{PC} = (V_{pc}, E_{pc})$, where $V_{pc}$ denotes a set of patients and cell lines as nodes and $E_{pc}$ denotes a set of paired patient-cell line edges. We utilized the *CTDPathSim2.0*[25] package to compute edge weights. *CTDPathSim2.0* integrated DNA methylation, gene expression, and CNA datasets from patient tumor samples and cell lines applying a deconvolution-based method followed by a biological pathway activity-based approach to compute the similarity scores between 0 to 1 with minmax normalization.

**Cell line-drug bipartite network.** The cell line-drug bipartite network ($\boldsymbol{B_{CP}}$) was built with cell line-drug similarity networks. $\boldsymbol{B_{CD}}$ can be defined as an undirected graph, $B_{CD} = (V_{cd}, E_{cd})$, where $V_{cd}$ denotes a set of cell lines and drugs as nodes and $E_{cd}$ denotes a set of paired cell line-drug edges. We retrieved log normalized $IC_{50}$ values of 397 drugs tested on 936 cell lines from CCLE and GDSC where $\ln(IC_{50}) < -2.0$ was used to define sensitive drug response. All other $\ln(IC_{50})$ values were considered as resistant drug responses. This threshold was chosen based on previous *in vitro* studies [26,27]. An edge weight was 1 if a cell line was sensitive to the drug; otherwise, resistant or unknown was set to 0.

**Drug-gene bipartite network.** The drug-gene bipartite network ($\boldsymbol{B_{DG}}$) was built with drug-gene interaction networks. $\boldsymbol{B_{DG}}$ can be defined as an undirected graph, $\boldsymbol{B_{DG}} = (V_{dg}, E_{dg})$, where $V_{dg}$ denotes a set of drugs and genes as nodes and $E_{dg}$ denotes a set of paired drug-gene edges. Databases such as the Comparative Toxicogenomics Database (CTD)[28], Therapeutic Target Database (TTD)[29], Guide to PHARMACOLOGY (GtoPdb)[30], and GDSC have screened at the large scale of the thousands of interactions between genes and drugs revealing the effectiveness and side effects of the drug therapy. We retrieved 263 drugs and 3,757 genes with 18,369 documented associations from these databases. An edge was assigned between a drug and a gene if an interaction was known; otherwise, no edge was assigned.

**Patient-gene bipartite network.** The patient-gene bipartite network ($\boldsymbol{B_{PG}}$) was built with patient-gene mutation edges. $\boldsymbol{B_{PC}}$ can be defined as an undirected graph, $\boldsymbol{B_{PG}} = (V_{pg}, E_{pg})$, where $V_{pg}$ denotes a set of patients and genes as nodes and $E_{pg}$ denotes a set of paired patient-gene edges. To build this network, we downloaded mutation data for tumor samples from the TCGA database. We filtered out the silent mutations and used all other mutations of 3,734 genes among 973 patients with 21,019 edges. An edge was set 1 if a gene was mutated in a patient.

**Patient-drug bipartite network.** The patient-drug bipartite network ($\boldsymbol{B_{PD}}$) was built with patient-drug response edges. $\boldsymbol{B_{PD}}$ can be defined as an undirected graph, $\boldsymbol{B_{PD}} = (V_{pd}, E_{pd})$, where $V_{pd}$ denotes a set of patients and drugs as nodes and $E_{pd}$ denotes a set of patient-drug edges. We prepared this network using drug response data for tumor samples from the TCGA database and processed it using OS and PFI as clinical endpoints following our previous works[25,31]. We also considered the time intervals in which drugs were applied. We considered survival status "censored" as the representative of sensitive drug response, whereas survival status "dead" was representative of resistant drug response as we mapped patients' survival status to drug response[25,31]. We assigned an edge between a cell line and a drug if the cell line was sensitive to that drug (i.e., edge weight = 1); otherwise, if the response was resistant or unknown, no edge was assigned (i.e., edge weight = 0). **Supplementary Table S2** shows the number of sensitive and resistant cases of the retrieved drugs that were applied in the TCGA BRCA sample.

**Supplementary Table S3** shows the final numbers of different nodes and edges in the multiplex and bipartite networks of *PDDRNet-MH* with breast cancer samples in the patient network.

**Supplementary Table S2: Number of sensitive and resistant breast cancer patients in TCGA.**

| Drug | Sensitive patient | Resistant patient |
|---|---|---|
| 5−Fluorouracil | 91 | 11 |
| Cyclophosphamide | 454 | 53 |
| Docetaxel | 175 | 18 |
| Doxorubicin | 317 | 41 |
| Epirubicin | 40 | 3 |
| Gemcitabine | 4 | 5 |
| Methotrexate | 22 | 6 |
| Paclitaxel | 195 | 30 |

| | | |
|---|---|---|
| Tamoxifen | 228 | 17 |
| Vinorelbine | 3 | 4 |
| Zoledronate | 15 | 4 |
| Cisplatin | 1 | 1 |
| Fulvestrant | 1 | 6 |
| Lapatinib | 2 | 1 |
| Mitomycin-C | 1 | 1 |
| Mitoxantrone | 2 | 0 |
| Pemetrexed | 1 | 0 |
| Vinblastine | 0 | 0 |
| Vincristine | 2 | 0 |

**Supplementary Table S3: Number of different nodes and edges in *PDDRNet-MH* with breast cancer samples.** n1 and n2 denote the number of two different node types in each bipartite network.

| | Network | Nodes | | Edges |
|---|---|---|---|---|
| **Multiplex** | Patient | 1,064 | | 1,204,506 |
| | Cell line | 715 | | 510,510 |
| | Drug | 397 | | 157,212 |
| | Gene | 4,313 | | 18,597,656 |
| | **n1-n2** | **n1** | **n2** | |
| **Bipartite** | Patient-cell line | 1,071 | 691 | 980,973 |
| | Cell line-drug | 658 | 397 | 263,140 |
| | Drug-gene | 263 | 3,757 | 18,369 |
| | Gene-patient | 3,734 | 973 | 21,019 |
| | Patient-drug (sensitive) | 577 | 17 | 1,552 |

**Running PDDRNet-MH**

After the construction of all the multiplex and bipartite networks, *PDDRNet-MH* includes two steps: (1) computing the global transition matrix and (2) performing RWR on the multiplex heterogeneous network.

Computing the global transition matrix of PDDRNet-MH:

Let's consider, the numbers of patients, cell lines, genes, and drugs in the multiplex heterogeneous network are *p*, *c*, *g,* and *d*, respectively. Following a previous approach[32], we define $A_{PP(p\times p)}$, $A_{CC(c\times c)}$, $A_{GG(g\times g)}$ and $A_{DD(d\times d)}$ as the adjacency matrices of $\boldsymbol{M_{PP}}$, $\boldsymbol{M_{CC}}$, $\boldsymbol{M_{GG}}$ and $\boldsymbol{M_{DD}}$, respectively; $B_{PC(p\times c)}$, $B_{CD(c\times d)}$, $B_{DG(d\times g)}$, $B_{PG(p\times g)}$ and $B_{PD(p\times d)}$ as the adjacency matrices of $\boldsymbol{B_{PC}}$, $\boldsymbol{B_{CD}}$, $\boldsymbol{B_{DG}}$, $\boldsymbol{B_{PG}}$ and $\boldsymbol{B_{PD}}$. For the ease of discussion, we show the construction of an adjacency matrix *A* corresponds to a generic multiplex network $\boldsymbol{M}$ and an adjacency matrix *B* for a generic bipartite network $\boldsymbol{B}$.

Suppose multiplex network $\boldsymbol{M}$ is a collection of *L* undirected graphs, i.e., layers sharing the same set of *n* nodes (note: for *PDDRNet-MH*, *L* = 4). Each layer $\alpha = 1, \ldots, L$, is defined by $n \times n$ adjacency matrix as:

$$A^{[\alpha]} = (A^{[\alpha]}(i,j))_{i,j=1,\dots,n} \quad (1)$$

where $A^{[\alpha]}(i,j) = 1$ if node *i* and *j* are connected on a layer $\alpha$, and 0 otherwise. Also, $A^{[\alpha]}(i,i) = 0$, i.e., no interaction between the same nodes. Multiplex network ***M*** is characterized by the adjacency matrix $A = A^{[1]},\dots, A^{[L]}$. Matrix *A* contains the different types of transitions defined as:

$$A = \begin{bmatrix} (1-\delta)A^{[1]} & \frac{\delta}{(L-1)}\mathbf{I} & \cdots & \frac{\delta}{(L-1)}\mathbf{I} \\ \frac{\delta}{(L-1)}\mathbf{I} & (1-\delta)A^{[2]} & \cdots & \frac{\delta}{(L-1)}\mathbf{I} \\ \vdots & \vdots & \ddots & \vdots \\ \frac{\delta}{(L-1)}\mathbf{I} & \frac{\delta}{(L-1)}\mathbf{I} & \cdots & (1-\delta)A^{[L]} \end{bmatrix} \quad (2)$$

where **I** represents the $n \times n$ identity matrix. The parameter $\delta \in (0,1)$ quantifies the surfer's probability of staying in a layer or jumping between the layers: if $\delta = 0$, the surfer will always stay in the same layer after a non-restart step. The diagonals of matrix *A* represent the potential intra-layer walks, whereas the off-diagonal elements account for the possible jumps between different layers in the same multiplex network.

We construct the adjacency matrix of bipartite network ***B*** following the previous work[33]. Let us consider a multiplex network, ***N*** composed of *n* nodes and *L*-layers with an adjacency matrix $A_{NN(n\times n)}$, like the one described in equation (2). Let also consider another multiplex network, ***M*** composed of *m* nodes and *L*-layers with an adjacency matrix $A_{MM(m\times m)}$. The bipartite graphs with adjacency matrices $B_{n\times m}^{1,\dots,L}$ associate the *n* and *m* nodes in each layer of the multiplex network ***N*** to each layer of the multiplex network ***M***. These bipartite graphs are identical, we define them as $B_{(n\times m)}$, and construct the bipartite adjacency matrix of the multiplex heterogeneous graph by sticking $B_{(n\times m)}$ $L\times L$ times:

$$B_{NM} = \begin{pmatrix} B_{(n\times m)} \\ B_{(n\times m)} \\ \vdots \\ B_{(n\times m)} \end{pmatrix} \quad (3)$$

Using Eq. (2) and Eq. (3), we constructed all the adjacency matrices of the multiplex networks and bipartite networks. Therefore, an adjacency matrix of the multiplex heterogeneous network of *PDDRNet-MH* can be represented as:

$$H = \begin{bmatrix} A_{PP} & B_{PC} & B_{PD} & B_{PG} \\ B_{PC}^T & A_{CC} & B_{CD} & 0 \\ B_{PD}^T & B_{CD}^T & A_{DD} & B_{DG} \\ B_{PG}^T & 0 & B_{DG}^T & A_{GG} \end{bmatrix} \quad (4)$$

where $B_{PC}^T$, $B_{CD}^T$, $B_{DG}^T$, $B_{PG}^T$ and $B_{PD}^T$ are the transpose matrices of $B_{PC}$, $B_{CD}$, $B_{DG}$, $B_{PG}$ and $B_{PD}$, respectively. In the next step, we compute the different transition probabilities of the random walk with restart on the multiplex heterogeneous network.

Let, $G = \begin{bmatrix} G_{PP} & G_{PC} & G_{PD} & G_{PG} \\ G_{CP} & G_{CC} & G_{CD} & 0 \\ G_{DP} & G_{DC} & G_{DD} & G_{DG} \\ G_{GP} & 0 & G_{GD} & G_{GG} \end{bmatrix}$ denotes the matrix of transition probabilities of the multiplex heterogeneous network (derived from the adjacency matrices in Eq. (4)). The diagonals describe the walks within a multiplex network, and off-diagonals elements describe the jumps between the multiplex networks. The computation of $G_{PP}$, $G_{PC}$, $G_{PD}$ and $G_{PG}$ (the first row of *G*) are shown below.

Suppose a surfer is at a patient node $p_i \in V_p$ in the patient multiplex network. In the next step, the surfer can either walk to a patient $p_j \in V_p$ with the transition probability:

$$G_{PP}(i,j) = \begin{cases} \frac{A_{PP}(i,j)}{\sum_{k=1}^{p} A_{PP}(i,k)}, & if \sum_{k=1}^{c} B_{PC}(i,k) = 0; \sum_{k=1}^{d} B_{PD}(i,k) = 0; \sum_{k=1}^{g} B_{PG}(i,k) = 0 \\ (1 - \frac{c\lambda}{b}) \frac{A_{PP}(i,j)}{\sum_{k=1}^{p} A_{PP}(i,k)}, & otherwise \end{cases} \quad (5)$$

where, λ denotes the inter-network jump probability, b denotes the maximum number of bipartite connections of the multiplex network and c denotes the count of all possible bipartite connections of $p_i$ in the same multiplex network with the nodes of other multiplex networks. Also, the surfer can jump from a patient node $p_i \in V_p$ through a bipartite network to other three multiplex networks using the following three equations:

(i) jump to a node $c_j \in V_c$ in the cell line network using bipartite connection with a transition probability:

$$G_{PC}(i,j) = \begin{cases} \lambda \frac{B_{PC}(i,j)}{\sum_{k=1}^{c} B_{PC}(i,k)}, & if \sum_{k=1}^{c} B_{PC}(i,k) \neq 0 \\ 0, & if \sum_{k=1}^{c} B_{PC}(i,k) = 0 \end{cases} \quad (6a)$$

(ii) jump to a node $d_j \in V_d$ in the drug network using bipartite connection with a probability:

$$G_{PD}(i,j) = \begin{cases} \lambda \frac{B_{PD}(i,j)}{\sum_{k=1}^{d} B_{PD}(i,k)}, & if \sum_{k=1}^{d} B_{PD}(i,k) \neq 0 \\ 0, & if \sum_{k=1}^{d} B_{PD}(i,k) = 0 \end{cases} \quad (6b)$$

(iii) jump to a node $g_j \in V_g$ in the gene network using bipartite connection with a probability

$$G_{PG}(i,j) = \begin{cases} \lambda \frac{B_{PG}(i,j)}{\sum_{k=1}^{g} B_{PG}(i,k)}, & if \sum_{k=1}^{g} B_{PG}(i,k) \neq 0 \\ 0, & if \sum_{k=1}^{g} B_{PG}(i,k) = 0 \end{cases} \quad (6c)$$

The other elements of the global transition matrix can be calculated in a similar way using appropriate adjacency matrices (i.e., elements of H) from Eq. 4.

**Effect of the multiplex networks in drug response prediction**

We built *PDDRNet-MH* with four multiplex networks. Here, the patient and drug multiplex networks were mandatory and cell line and gene multiplex networks were optional, i.e., the tool can be run without these multiplex networks. We checked the effect of cell line and gene multiplex networks in the prediction performance of *PDDRNet-MH.* For this purpose, we performed three sets of LOOCV experiments (see **Materials and Methods**) for each of the eleven drugs by (1) removing cell line multiplex network, (2) removing gene multiplex network, and (3) removing both cell line and gene multiplex networks. We present the AUC values of these runs in **Supplementary Table S4**. Removing the cell line multiplex network from *PDDRNet-MH* decreased the ROC-AUC and PR-AUC values in 80% of the drugs. On the other hand, removing both cell line and gene multiplex networks decreased ROC-AUC and PR-AUC values in almost 60% of the total drugs, and removing only gene multiplex network decreased ROC-AUC and PR-AUC values in nearly 40% of the total drugs. This experiment suggests that the cell line multiplex network was an important network to achieve higher AUC in *PDDRNet-MH.* The gene network can also affect the prediction performance by lowering the AUCs. Interestingly, for the two drugs, such as 5-fluorouracil and epirubicin, removing both cell line and gene multiplex networks slightly improved prediction performance. Overall, *PDDRNet-MH* consistently produced high AUC values for both the ROC and PR curves with all four multiplex networks.

**Supplementary Table S4: ROC-AUC and PR-AUC values of the eleven drugs for TCGA BRCA samples in checking the impact of multiplex networks.** * and †, show the higher and lower values compared to *PDDRNet-MH* with all four multiplex networks, respectively.

| **Drug** | **AUC-C**[a] | | **AUC-G**[b] | | **AUC-CG**[c] | |
|---|---|---|---|---|---|---|
| | ROC | PR | ROC | PR | ROC | PR |
| 5−Fluorouracil | 0.54† | 0.89† | 0.61* | 0.91 | 0.62* | 0.93* |

| | | | | | | |
|---|---|---|---|---|---|---|
| Cyclophosphamide | 0.63† | 0.93† | 0.65 | 0.94 | 0.65 | 0.94 |
| Docetaxel | 0.73† | 0.96† | 0.74† | 0.97 | 0.72† | 0.96† |
| Doxorubicin | 0.64† | 0.93 | 0.64† | 0.92† | 0.64† | 0.93 |
| Epirubicin | 0.48† | 0.93† | 0.88* | 0.99* | 0.79* | 0.98* |
| Gemcitabine | 1.00 | 1.00 | 1.00 | 1.00 | 1.00 | 1.00 |
| Methotrexate | 0.88† | 0.96† | 0.95 | 0.99 | 0.89† | 0.97† |
| Paclitaxel | 0.69* | 0.97* | 0.61† | 0.90 | 0.62† | 0.91* |
| Tamoxifen | 1.00* | 1.00* | 0.64† | 0.96† | 0.65† | 0.96† |
| Vinorelbine | 0.82† | 0.95† | 1.00 | 1.00 | 1.00 | 1.00 |
| Zoledronate | 0.54† | 0.89† | 0.92† | 0.98† | 0.78† | 0.94† |

*Note:* [a]*Removing only cell line network;* [b]*removing only gene network;* [c]*removing both cell line and gene networks.*

**Layers of the patient multiplex network had the most considerable contribution to the drug response prediction**

In *PDDRNet-MH*, we used four parameters such as $\tau_p$, $\tau_c$, $\tau_d$ and $\tau_g$ as restart probabilities for multiplex networks, $\boldsymbol{M_{PP}}$, $\boldsymbol{M_{CC}}$, $\boldsymbol{M_{DD}}$ and $\boldsymbol{M_{GG}}$, respectively. These parameters control the probability of restart in the different layers of the corresponding multiplex network and can be explored to determine the importance of the layers in each multiplex network. For instance, we could favor the restart of the surfer in *Layer 1* (i.e., gene expression) of the patient multiplex network and hinder it in *Layer 4* (i.e., clinical similarity) to check the effect of these layers in AUC values. **Supplementary Table S5** does not show notable differences in the performances of the LOOCV experiment associated with the variations of the parameters $\tau_c$, $\tau_d$ and $\tau_g$ of the eleven drugs in terms of AUC values. Alteration in these parameters did not decrease the overall performance. However, the variations in $\tau_p$ altered the mean AUC values, showing that all the patient multiplex layers contributed to achieving high AUCs. Hence, equal weights should be applied to all these layers to achieve the highest AUC. Overall, in RWR, the surfer continued exploring the different network layers with the variation in these parameters and leveraging the combined network information, even if it did not restart in the seeds of one of the layers of *PDDRNet-MH*.

**Supplementary Table S5: Mean AUC with a standard deviation of ROC and PR curves of the eleven drugs for TCGA BRCA samples in checking the impact of the restart parameters associated with multiplex networks.** The parameter values in each multiplex network are in (*Layer1*, *Layer2, Layer3, Layer4*)/L format, where L = 4; * shows the default values for running *PDDRNet-MH*.

| Values in layers | ROC | | | | PR | | | |
|---|---|---|---|---|---|---|---|---|
| | $\tau_p$ | $\tau_c$ | $\tau_d$ | $\tau_g$ | $\tau_p$ | $\tau_c$ | $\tau_d$ | $\tau_g$ |
| **(.1,1.9,1,1)/L** | 0.55 ± 0.42 | 0.77 ± 0.17 | 0.78 ± 0.16 | 0.77 ± 0.17 | 0.85 ± 0.23 | 0.96 ± 0.04 | 0.96 ± 0.03 | 0.96 ± 0.04 |
| **(1,.1,1.9,1)/L** | 0.54 ± 0.43 | 0.77 ± 0.17 | 0.77 ± 0.17 | 0.77 ± 0.17 | 0.84 ± 0.22 | 0.96 ± 0.04 | 0.96 ± 0.04 | 0.96 ± 0.04 |
| **(1,1,.1,1.9)/L** | 0.54 ± 0.41 | 0.77 ± 0.17 | 0.77 ± 0.17 | 0.77 ± 0.17 | 0.84 ± 0.22 | 0.96 ± 0.04 | 0.96 ± 0.04 | 0.96 ± 0.04 |
| **(1.9,1,1,.1)/L** | 0.54 ± 0.42 | 0.77 ± 0.17 | 0.76 ± 0.17 | 0.77 ± 0.17 | 0.85 ± 0.22 | 0.96 ± 0.04 | 0.96 ± 0.04 | 0.96 ± 0.04 |

| | | | | | | | | |
|---|---|---|---|---|---|---|---|---|
| **(0,0,0,4)/L** | 0.48 ± 0.50 | 0.77 ± 0.17 | 0.78 ± 0.16 | 0.77 ± 0.17 | 0.83 ± 0.23 | 0.96 ± 0.04 | 0.96 ± 0.04 | 0.96 ± 0.04 |
| **(0,0,4,0)/L** | 0.49 ± 0.49 | 0.77 ± 0.17 | 0.78 ± 0.16 | 0.77 ± 0.17 | 0.83 ± 0.23 | 0.96 ± 0.04 | 0.96 ± 0.03 | 0.96 ± 0.04 |
| **(0,4,0,0)/L** | 0.50 ± 0.49 | 0.77 ± 0.17 | 0.78 ± 0.16 | 0.77 ± 0.17 | 0.83 ± 0.23 | 0.96 ± 0.04 | 0.96 ± 0.03 | 0.96 ± 0.04 |
| **(4,0,0,0)/L** | 0.49 ± 0.49 | 0.77 ± 0.17 | 0.76 ± 0.18 | 0.77 ± 0.17 | 0.83 ± 0.23 | 0.96 ± 0.04 | 0.96 ± 0.04 | 0.96 ± 0.04 |
| **(1,1,1,1)/L*** | 0.77 ± 0.17 | 0.77 ± 0.17 | 0.77 ± 0.17 | 0.77 ± 0.17 | 0.96 ± 0.04 | 0.96 ± 0.04 | 0.96 ± 0.04 | 0.96 ± 0.04 |

### ***PDDRNet-MH*** **was robust with moderate changes in the parameters**

We checked the influence of the parameters such as $r$, δ, $\eta_p$, and $\eta_d$ involved in the *PDDRNet-MH* using the LOOCV strategy proposed in Materials and Methods for the eleven drugs. Here, $r$ was the global restart parameter for which the default value was set as $r = 0.7$, following the earlier publications[32–34]. $r$ controls the probability of jumping back to the seed nodes during the random walk. **Supplementary Figure S1** shows the effects of different $r$ values on the AUC of ROC and PR curves for randomly chosen three drugs for this demonstration. As the value of $r$ increased, the performance got better. Larger $r$ values caused RWR not to diffuse the information to farther distances; instead, it maintained the diffusion to be close to the seed nodes. Overall, as the value of $r$ increased from 0.1 to 0.5, the AUC values improved significantly (p-value < 0.05) for the eleven drugs (**Supplementary Table S5**). The increase in $r$ from 0.5 to 0.7 and further, 0.9 did not change the overall performance (p-value > 0.05) of *PDDRNet-MH*.

We then studied the effect of hyperparameters related to the random walks in multiplex networks, such as δ. The default value of δ was set as 0.5. δ quantifies the probability that the surfer jumps from the current node to the same node in a different layer after a non-restart step. If δ = 0, the surfer would always stay in the same layer, and if δ = 1, the surfer would jump to a different layer at each step. We did not observe notable changes in the AUC values of PR curves for the eleven drugs with variations in this parameter (**Supplementary Table S5**).

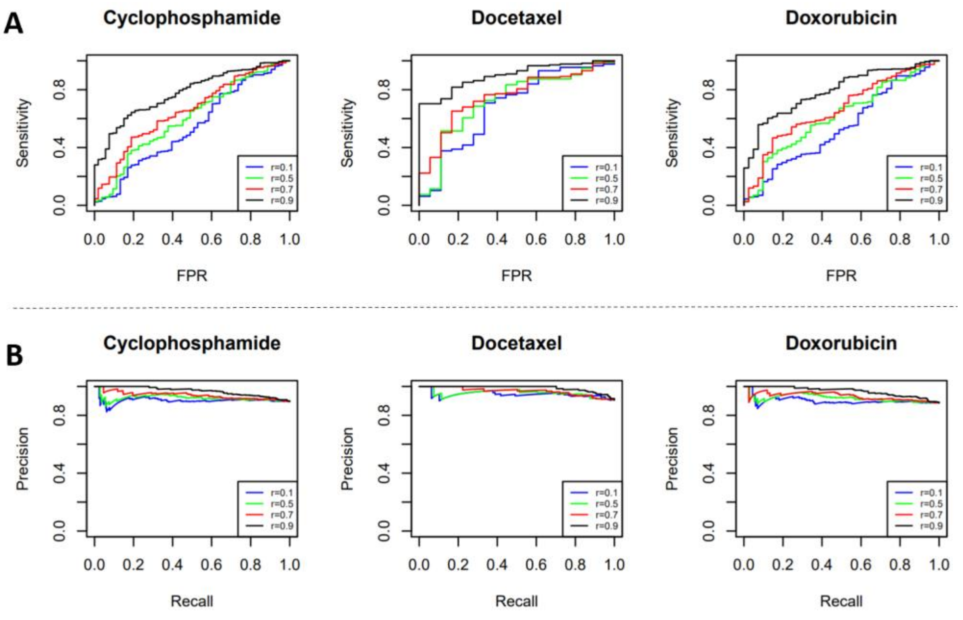


**Supplementary Figure 1. A) AUC-ROC and B) AUC-PR curves for the three drugs with variation in global restart parameter *r*.**

We checked the performance of the hyperparameters $\eta_p$ and $\eta_d$. Here, $\eta_p$ and $\eta_d$ quantify the probability of restart of the seeds in the patient multiplex and the drug multiplex networks, respectively. For these, the default values were, $\eta_p = 0.75, \eta_d = 0.25$. High value of $\eta_p$ means high likelihood to restart the walk from a sensitive patient seed. The high value of $\eta_p$ (≥ 0.5) produced high AUC (**Supplementary Table S5**). In LOOCV, we had only one seed in the drug multiplex network and multiple seeds in the patient multiplex network (see **Materials and Methods**), hence low $\eta_p$ values decreased the AUC values.

**Supplementary Table S5: Mean AUC with a standard deviation of ROC and PR curves of the eleven drugs for TCGA BRCA samples in checking the impact of the parameters in PDDRNet-MH.** * shows the default values.

| | | | | | |
|---|---|---|---|---|---|
| **ROC** | | | | | |
| ***r* = 0.1** | 0.57 ± 0.11 | **δ = 0.1** | 0.76 ± 0.17 | **$\eta_p$= 0.1, $\eta_d$ = 0.9** | 0.32 ± 0.29 |
| ***r* = 0.5** | 0.69 ± 0.18 | **δ = 0.5*** | 0.77 ± 0.17 | **$\eta_p$= 0.5, $\eta_d$ = 0.5** | 0.75 ± 0.19 |
| ***r* = 0.7*** | 0.77 ± 0.17 | **δ = 0.9** | 0.75 ± 0.15 | **$\eta_p$= 0.75, $\eta_d$ = 0.25*** | 0.77 ± 0.17 |
| ***r* = 0.9** | 0.89 ± 0.11 | | | **$\eta_p$= 0.9, $\eta_d$ = 0.1** | 0.78 ± 0.16 |
| **PR** | | | | | |
| ***r* = 0.1** | 0.85 ± 0.10 | **δ = 0.1** | 0.96 ± 0.04 | **$\eta_p$= 0.1, $\eta_d$ = 0.9** | 0.75 ± 0.27 |
| ***r* = 0.5** | 0.93 ± 0.04 | **δ = 0.5*** | 0.96 ± 0.04 | **$\eta_p$= 0.5, $\eta_d$ = 0.5** | 0.95 ± 0.04 |
| ***r* = 0.7*** | 0.96 ± 0.04 | **δ = 0.9** | 0.95 ± 0.03 | **$\eta_p$= 0.75, $\eta_d$ = 0.25*** | 0.96 ± 0.04 |
| ***r* = 0.9** | 0.98 ± 0.02 | | | **$\eta_p$= 0.9, $\eta_d$ = 0.1** | 0.96 ± 0.03 |

Overall, *PDDRNet-MH* was a robust algorithm since moderate variations in the parameters did not lead to significant variations in the drug response prediction.

**Running state-of-the-art-methods**

We compared PDDRNet-MH with the two state-of-the-art machine learning methods, namely, IDWAS and SVM-RFE.

**Running IDWAS**

To run IDWAS, we used gene expression data (RNA-Seq) for TCGA samples in FPKM format and 1,019 CCLE cell lines in RPKM format with 30,681 common genes. We utilized the R package *pRRophetic*[35] for running IDWAS. *pRRophetic* applied a quantile normalization as a batch-correction method to the combined gene expression matrix of samples and cell lines and removed the 20% of genes with the lowest variability in expression across all the samples and cell lines. In the next step, a linear ridge regression model was employed with 10-fold cross-validation with the cell lines' gene expression. Here, the phenotype data as drug sensitivity measurements with power-transformed $IC_{50}$ values were used as response variables. After the model was fitted, it was then applied to the tumor sample's gene expression matrix, thus predicting each patient's drug response estimate. We applied this model for each drug used in the study. The estimated scores (E) were compatible with $IC_{50}$ values (means the lower the value better the score); hence we applied a sigmoid transformation (i.e., *Sig*(E)) on the estimated drug response values and used 1- *Sig*(E) as the final scores comparable with the RWR scores of *PDDRNet-MH.*

**Running SVM-RFE**

For running SVM-RFE, we utilized the gene expression datasets of TCGA samples and CCLE cell lines. For cell lines' drug response, $\ln(IC_{50}) < -2.0$ was used to define "sensitive" drug response and other $\ln(IC_{50})$ values were for the "resistant" drug response. We utilized cell lines' gene expression data in the SVM algorithm using the *Caret*[36] R package to develop a drug response model using two classes: drug-sensitive and drug-resistant. This model was then applied to patients' gene expression matrix to predict drug response-based classification. Following the SVM-RFE model[37], we applied the SVM algorithm utilizing a recursive feature elimination (RFE) strategy on cell lines' gene expression to select 500 important features (i.e., genes). Then another SVM model was employed on patients' gene expression matrix utilizing these RFE-selected features to classify the patient samples into drug-sensitive and resistant groups. Since SVM's output was binary, we used decision values as drug response prediction scores. We could not run SVM-RFE for the drugs if the cell lines had only one label, i.e., either sensitive or resistant.
